# Evolutionary dynamics of *Tomato spotted wilt virus* within and between alternate plant hosts and thrips

**DOI:** 10.1101/2020.01.13.904250

**Authors:** Casey L. Ruark-Seward, Brian Bonville, George Kennedy, David A. Rasmussen

## Abstract

Tomato spotted wilt virus (TSWV) is a generalist pathogen with one of the broadest known host ranges among RNA viruses. To understand how TSWV adapts to different hosts, we experimentally passaged viral populations between two alternate hosts, *Emilia sochifolia* and *Datura stramonium*, and an obligate vector in which it also replicates, western flower thrips (*Frankliniella occidentalis*). Deep sequencing viral populations at multiple time points allowed us to track the evolutionary dynamics of viral populations within and between hosts. High levels of viral genetic diversity were maintained in both plants and thrips between transmission events. Rapid fluctuations in the frequency of amino acid variants indicated strong host-specific selection pressures on proteins involved in viral movement (NSm) and replication (RdRp). While several genetic variants showed opposing fitness effects in different hosts, fitness effects were generally positively correlated between hosts indicating that positive rather than antagonistic pleiotropy is pervasive. These results suggest that high levels of genetic diversity together with the positive pleiotropic effects of mutations have allowed TSWV to rapidly adapt to new hosts and expand its host range.

## Introduction

Despite theoretical predictions that specialist pathogens should outcompete generalists, multi-host pathogens are abundant in nature (Woolhouse *et al.*, 2001). One extreme example of such generalism is provided by plant viruses, which unlike their animal infecting counterparts, often infect hundreds or even thousands of phylogenetically distant plant species (Dawson and Hilf, 1992; Moury *et al.*, 2017). In turn, generalist plant viruses are often transmitted by generalist plant-feeding insects such as aphids, thrips and whiteflies, which feed on a wide range of plants (Nault, 1997; Jones, 2005; Gilbertson *et al.*, 2015). Thus, highly polyphagous vectors routinely transmit viruses between different plant species, which may strongly favor generalists that can either readily adapt or remain adapted to a wide range of potential hosts.

Tomato spotted wilt virus (TSWV) is a negative-stranded RNA virus in the genus Orthotospovirus in the order *Bunyavirales* (Abudurexiti *et al.*, 2019). Even when considered among other plant viruses, TSWV is a generalist *par excellence*, with a described host range of over 1,000 plant species distributed across more than 90 families of angiosperms (Parrella, *et al.*, 2003; Oliver and Whitfield, 2016). Important hosts include Solanaceous crops like tomato, pepper and tobacco, of which TSWV is a major constraint on production worldwide (Pappu *et al.*, 2009). Tospoviruses like TSWV also have the rather rare ability among plant viruses to persist and replicate in their insect vector, thrips (Whitfield *et al.*, 2005). While little is known about the realized host range of any particular genotype of TSWV in nature, TSWV vectors like western flower thrips (*Frankliniella occidentalis*) feed on hundreds of plant species (Jones, 2005), making it highly probable that a single viral lineage may move between multiple crop and wild host species over the course of a single growing season and then overwinter in another perennial host (Groves *et al.*, 2002).

What factors ultimately shape TSWV’s broad host range and how the virus adapts to novel hosts remains little explored. Among RNA viruses more generally, fitness tradeoffs between alternative hosts are widely assumed to limit simultaneous adaptation to multiple hosts (Elena *et al.*, 2009; García-Arenal and Fraile, 2013; Longdon *et al.*, 2014). Consistent with these predictions, experimental evolution studies in which viruses are passaged between alternate hosts and/or vectors have provided evidence that mutations that increase fitness in one host often decrease fitness in another host, suggesting that antagonistic pleiotropy underlies fitness tradeoffs between hosts (Duffy *et al.*, 2006; Lalic *et al.*, 2011). Nevertheless, generalists with high fitness across multiple hosts often evolve in experimental evolution studies where microbes are serially passaged between different host environments (Weaver *et al.*, 1999; Kassen, 2002; Bedhomme *et al.*, 2012). Thus, the extent to which fitness tradeoffs actually limit adaptation to multiple hosts remains unclear, especially for generalist pathogens like TSWV whose past ecological success suggests that the virus may have evolved efficient strategies to circumvent fitness tradeoffs and readily adapt to new host environments.

To address these questions, we experimentally passaged a field-collected isolate of TSWV between plants using western flower thrips as a vector (Figure 1). In one line, we alternately passaged the virus between two plant species: *Emilia sochifolia* (Asteraceae) and *Datura stramonium* (Solanaceae). In two other lines, we passaged the virus exclusively on either *Emilia* or *Datura*. Hereafter, we refer to these as the *Alternating, Emilia* and *Datura* lines. During each passage cycle, the viral population was deep sequenced at multiple time points in plants and once in thrips. This time-resolved deep sequencing data allowed us to track the evolutionary dynamics of viral populations both within and between hosts, and thereby quantify the fitness of genetic variants in different host environments.

**Figure 1:**
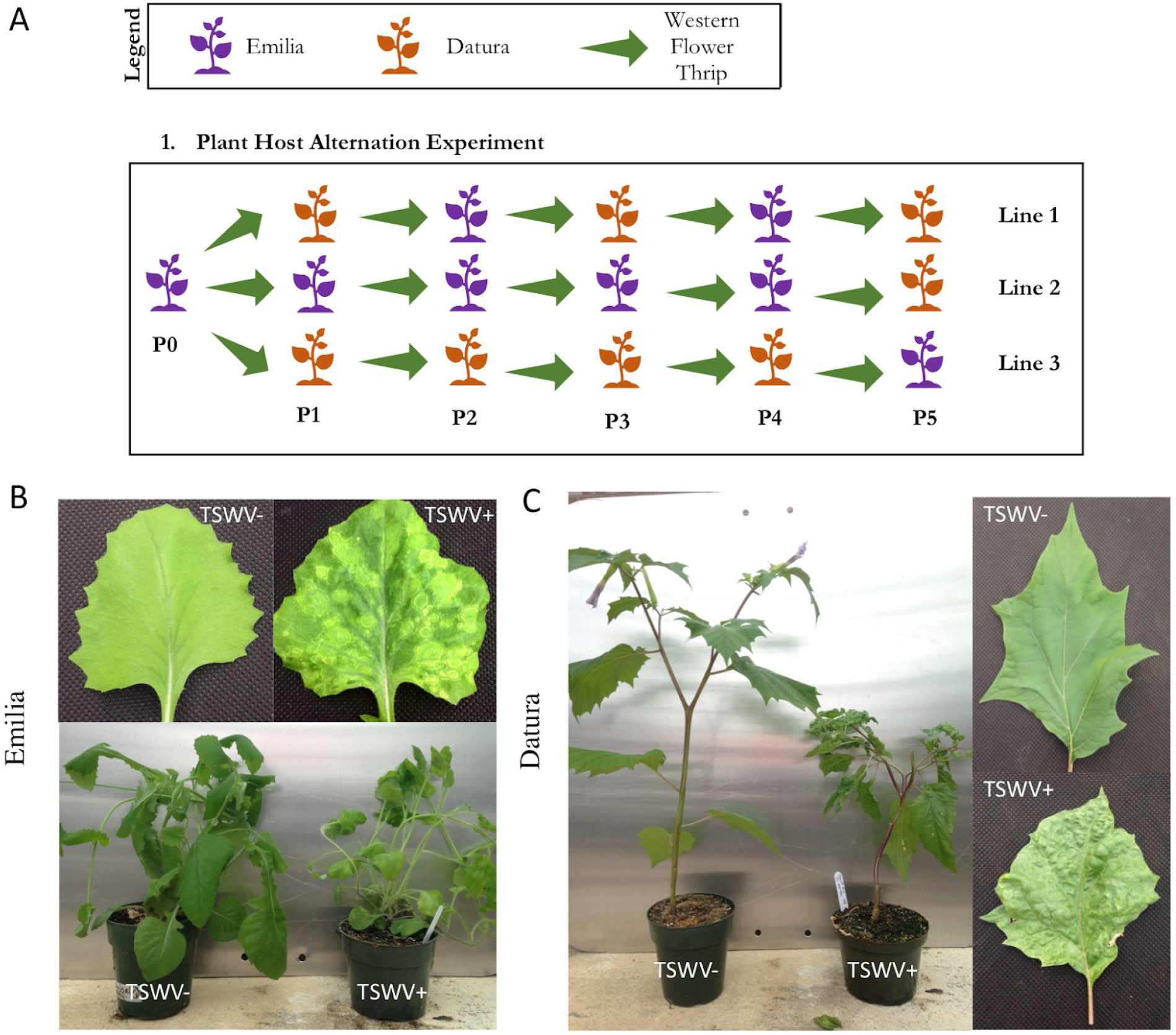
Schematic overview of the viral passaging experiment. **(A)** Virus was passaged for five generations alternately on *Emilia* and *Datura* (Line 1), passaged exclusively on *Emilia* and then back passaged to *Datura* (Line 2), or passaged exclusively on *Datura* and then back passaged to *Emilia* (Line 3). (B) Comparison of infected (TSWV+) and uninfected control (TSWV-) *Emilia* plants at passage P1. Infected *Emilia* leaves display the classic ringspot symptoms associated with TSWV. (C) Comparison of infected and uninfected *Datura* plants at passage P1.

## Results

### Deep sequencing of TSWV populations

Viral populations were deep sequenced twice at each sampling time point using paired sequencing replicates that originated from two independent reverse transcription reactions. Sequencing provided high coverage across all three segments of the TSWV genome, with a depth of coverage generally >1000X in both sequencing replicates (Supp. Figure 1). To ensure that genetic variants in the sequence reads represented actual variants present in the viral population, a conservative variant calling scheme was used in which a variant needed to be present at a frequency of at least 3% in both sequencing replicates in order to be called a true variant (see Methods). Called variants are therefore unlikely to represent RT, PCR or sequencing errors. Furthermore, single nucleotide variant (SNV) frequencies were highly correlated between the paired sequencing replicates (Figure 2), suggesting that our sequencing protocol introduced little sampling variance and that the sequence data accurately reflects the genetic composition of the original viral populations.

**Figure 2:**
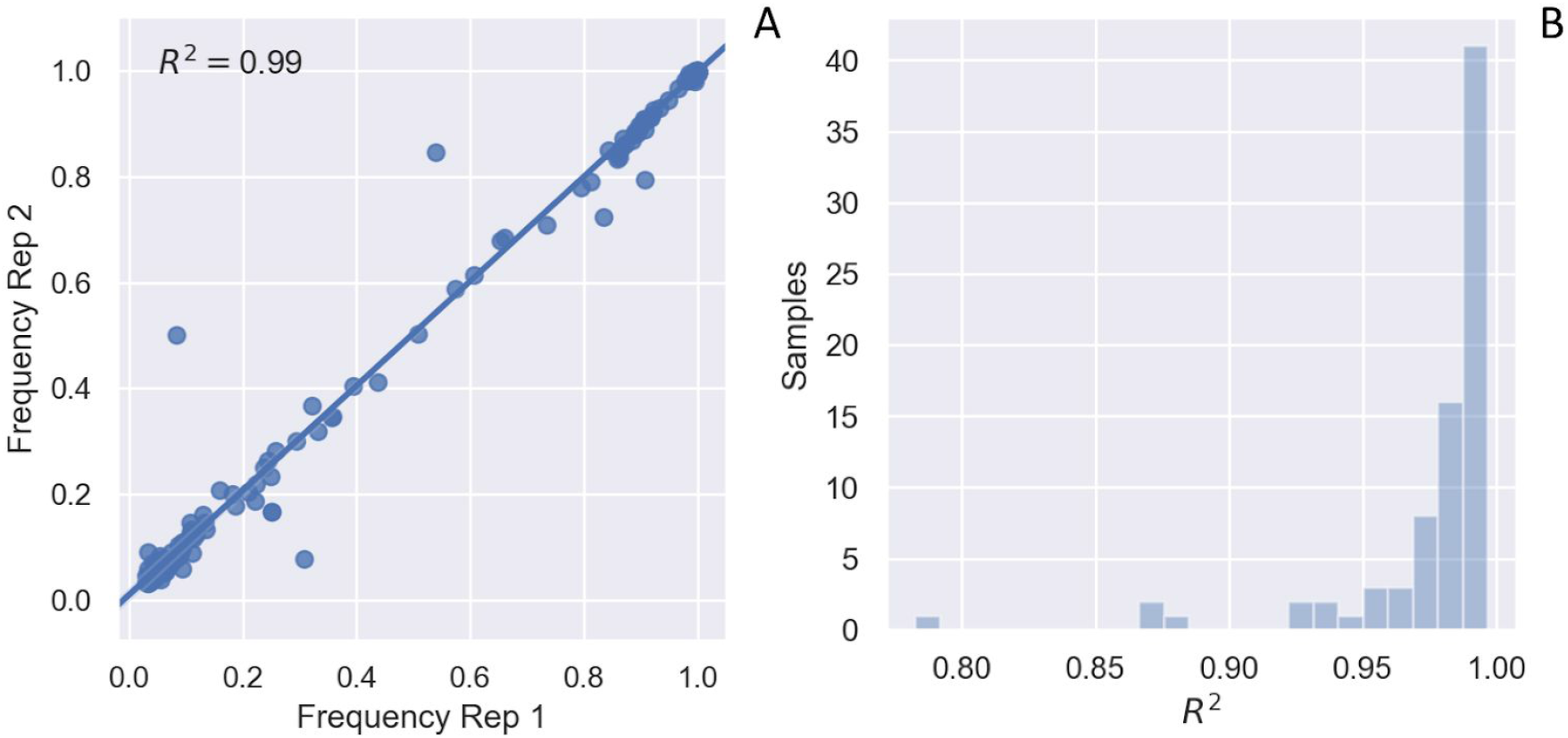
Single nucleotide variants (SNV) frequencies in paired sequencing replicates. **(A)** Frequency of individual SNVs at passage cycle P0 in two paired sequencing replicates obtained from independent reverse transcription (RT) reactions. The R^2^ value for variant frequencies between each paired replicate was determined by least-squares regression. **(B)** Histogram showing the distribution of R^2^ values between paired replicates at all time points sampled during the passaging experiment.

### Within and between host patterns of viral genetic diversity

First, the initial viral population in the field-collected tomato fruit (TF2) was sequenced. Clear hotspots of genetic diversity can be seen in the TF2 population, especially in intergenic regions which contained a large number of SNVs and indels (Figure 3). Protein coding regions exhibit less overall variability, but there is still considerable nonsynonymous variation. Of the 49 SNVs that fall within coding regions, 33 (67%) are predicted amino acid variants (AAVs) based on their translated sequences. There is also a hotspot of coding diversity at the 5’ end of the movement protein NSm. Viral diversity was similar in a leaf sampled from the same tomato plant in the field (Supp. Figure 2).

**Figure 3:**
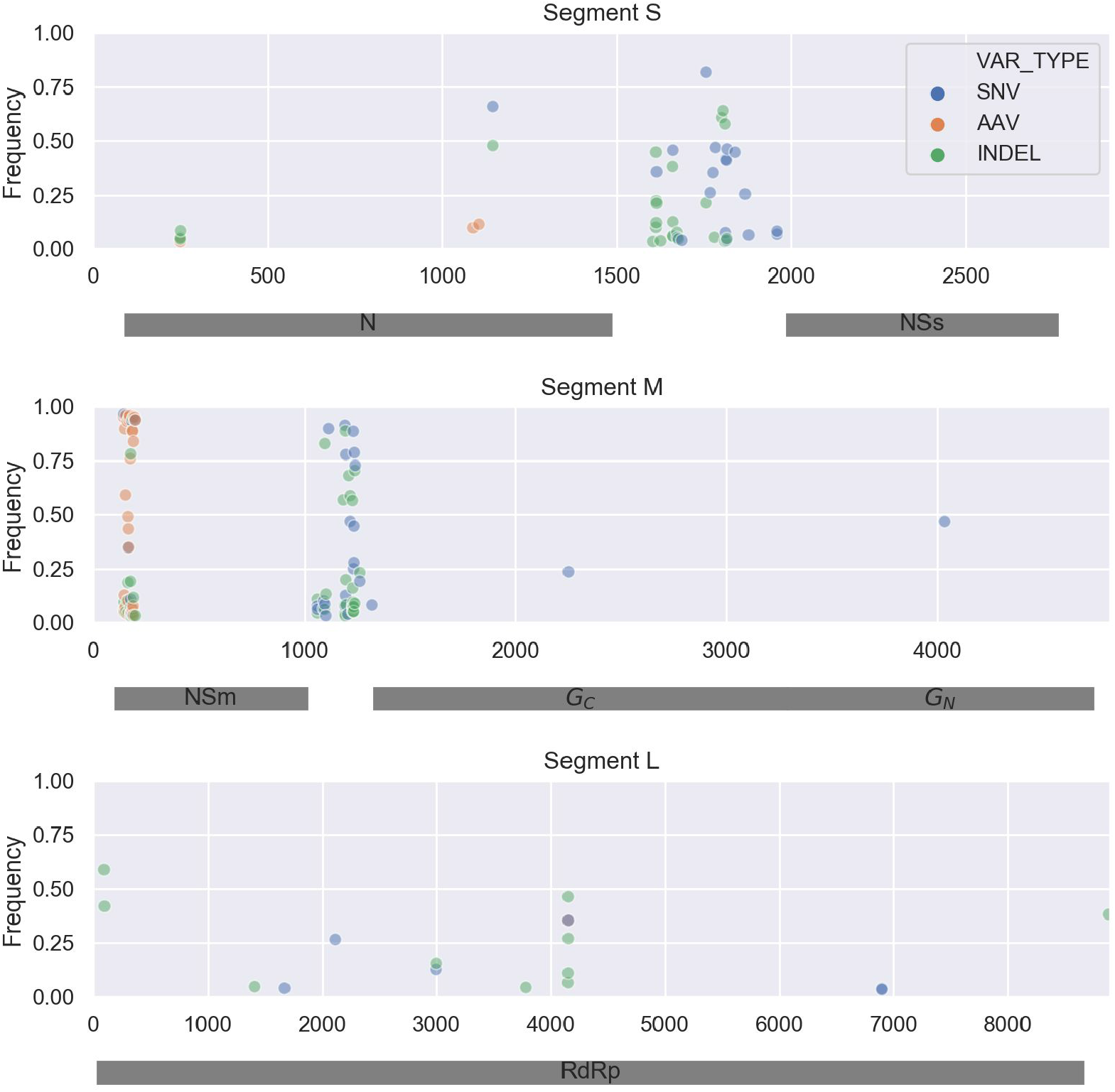
Viral genetic diversity in a naturally infected tomato fruit (TF2) collected in the field. All three passaging lines were derived from this TF2 isolate. The frequency of each single nucleotide variant (SNV), amino acid variant (AAV) and indel is plotted at its respective position along the three segments of TSWV’s genome. Shaded grey rectangles represent protein coding regions.

TSWV was mechanically transferred from the field collected tomato sample to a single *Emilia* plant, P0, from which all three lines were derived. In P0, the average pairwise distance between viral sequences was 25 single nucleotide substitutions, or 1.6 x 10^-3^ mutations per site. Average pairwise diversity tended to decrease slightly over time in all lines, with diversity declining more rapidly in *Datura* than in *Emilia* (Figure 4; blue). In contrast, genetic diversity tended to rebound whenever the virus was passaged back through thrips. No severe losses in diversity were observed when the virus was passaged from plants into thrips or from thrips back into plants, indicating a lack of severe population bottlenecks at transmission events. This may be an artefact of using multiple thrips to passage the virus between hosts and pooling these thrips for sequencing, but also may be due to TSWV’s ability to replicate in thrips. All three viral lines diverged from P0 by about 5-10 mutations by P5, but divergence appeared to slow after the first two passage cycles (Figure 4; orange).

**Figure 4:**
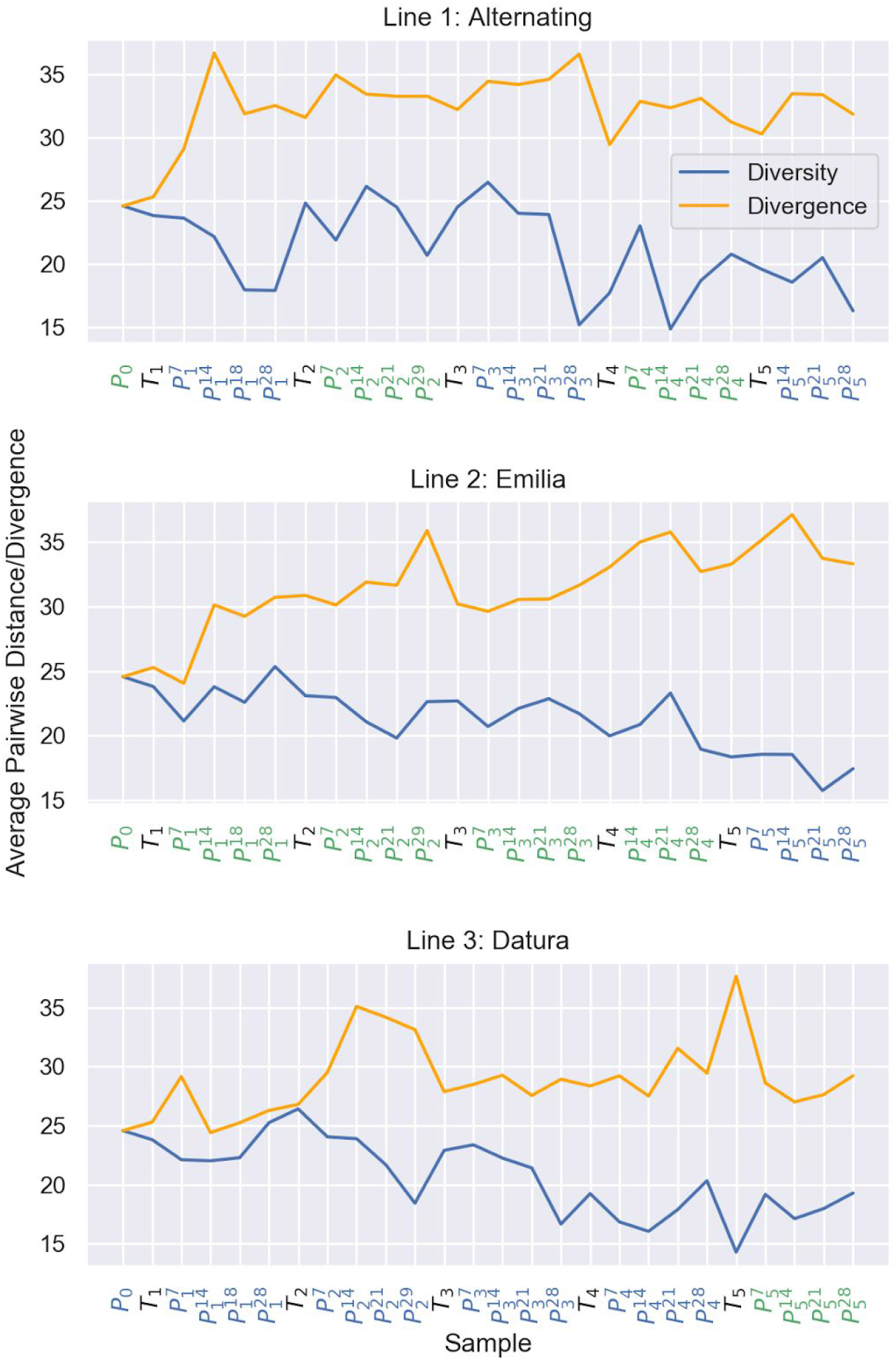
Genetic distance within and between viral populations at each sampled time point. The average pairwise distance within populations is shown in blue and the average pairwise divergence between each population and the founder viral population at P0 is shown in orange. Subscripts in the sample names denote passage number, superscripts denote days post infection. Samples from plants (P) are colored according to whether they were from *Emilia* (green) or *Datura* (blue). Samples from thrips (T) are colored black.

Within-host genetic diversity tended to mirror species-level diversity in TSWV samples collected around the world (Figure 5). Although within-host diversity is lower than species-level diversity, hotspots of diversity can be seen both in intergenic regions as well as in certain protein coding regions, especially at the N-terminus of NSm.

**Figure 5:**
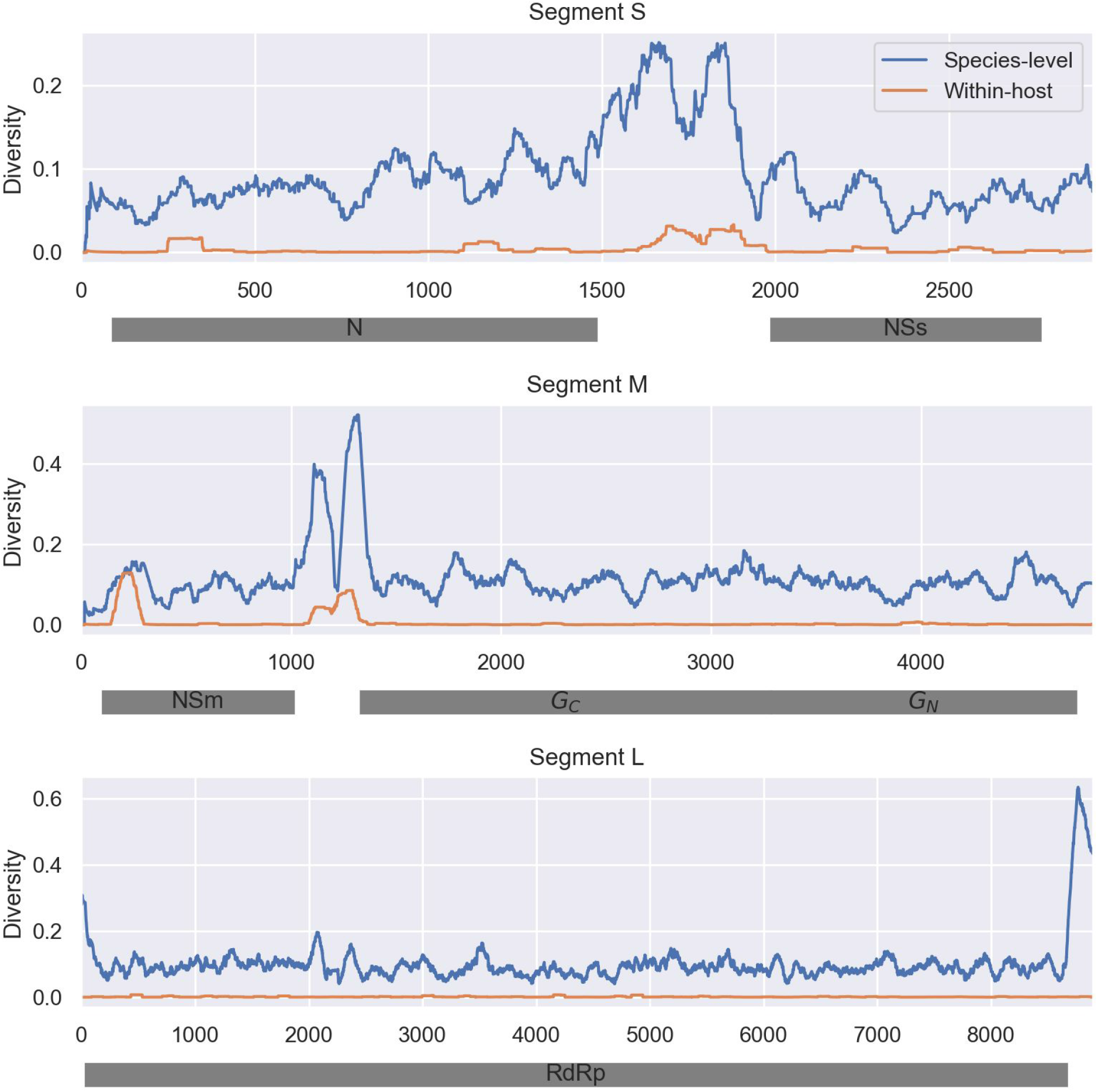
Comparison of within-host versus species-level genetic diversity in the TSWV genome. Genetic diversity is measured in terms of the entropy in nucleotide frequencies at each site in the genome. Within-host diversity values were averaged across all samples in the *Alternating* line. Species-level diversity was measured using a globally representative sample of publicly available TSWV samples (Supp. Table 1). Local fluctuations in diversity were smoothed by taking a running average across a 100 bp sliding window.

### Evolutionary dynamics of individual variants

While divergence between viral populations was low at the genome-wide level, tracking the evolutionary dynamics of individual variants revealed rapid and often host-specific changes in variant frequencies over time. The evolutionary dynamics of all SNVs in the *Alternating* line through time is shown in Supp. Figure 3. Because variants that are differentially selected between hosts are of particular interest, we looked for variants that were enriched in either plant host or the vector. Here, a variant is considered to be enriched if its average frequency was >5% higher in one host than in an alternate host, where the average is computed over all sampled time points.

To more easily visualize the evolutionary dynamics of individual variants, only amino acid variants (AAVs) are displayed for the *Alternating* line in Figure 6. Figure 6a shows the evolutionary dynamics of variants enriched in one plant host (*Emilia* or *Datura*) relative to the other. Variants can be seen that increase in frequency in *Emilia* but decline in *Datura* (e.g. NSm V17G) as well as variants that increase in frequency in *Datura* but decline in *Emilia* (e.g. NSm N22S). Figure 6b shows AAVs that are enriched in plants versus thrips or in thrips versus plants. There are quite a few variants on NSs, G_N_/G_C_ and RdRp that change dramatically in frequency whenever the virus is passaged between plants and thrips. Of the 15 variants that are enriched between plants and thrips, five variants are consistently enriched across all three experimental lines (Figure 7). Four of these five are enriched in plants versus thrips while only one variant (G_C_ E362D) was enriched in thrips versus plants.

**Figure 6:**
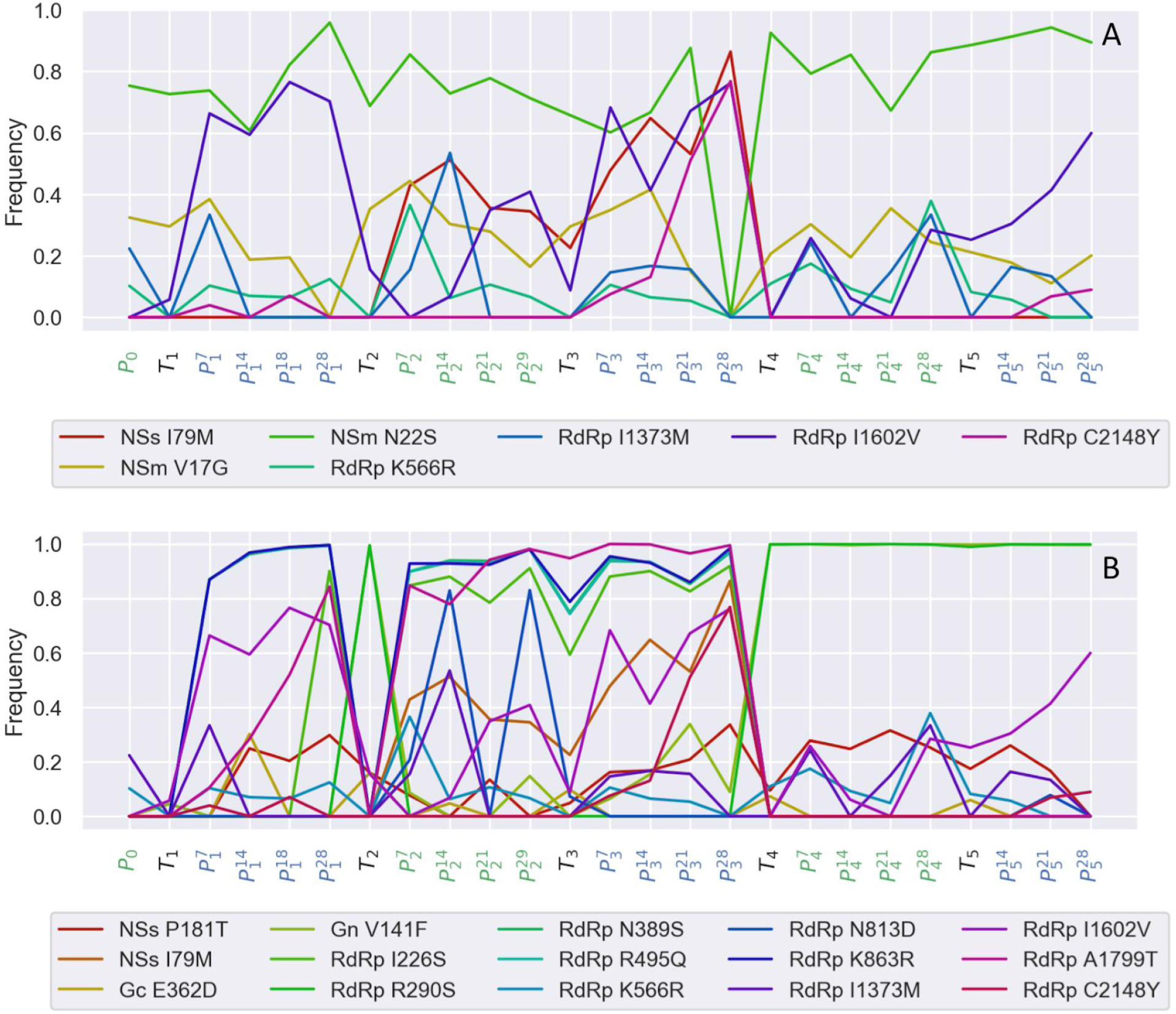
Evolutionary dynamics of individual amino acid variants in the *Alternating* line. **(A)** Time series of variants enriched in either *Emilia* or *Datura* relative to the other plant host. **(B)** Time series of variants enriched in either plants or thrips. A variant was considered to be enriched if its average frequency in one host was >5% higher than in the alternate host, where the average was computed over all sampled time points in each host. Sampling time points are colored by host; green = *Emilia*, black = thrips and blue = Datura.

**Figure 7:**
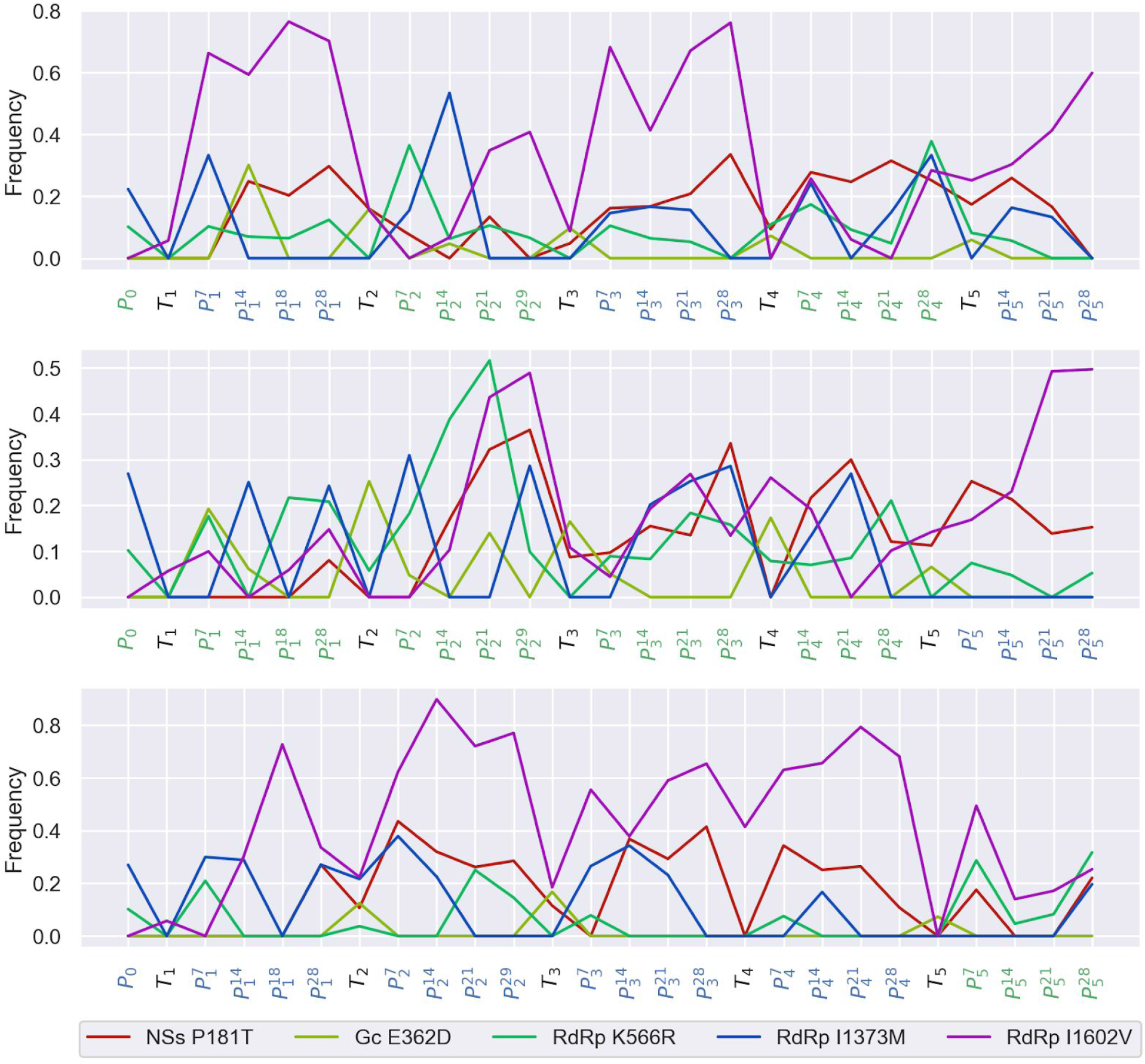
Evolutionary dynamics of individual amino acid variants consistently enriched in plants or thrips across all three lines. The three lines are: **(A)** *Alternating*, **(B)** *Emilia* and **(C)** *Datura*.

### Fitness effects between hosts and between plants and thrips

To more precisely quantify the fitness effects of variants in different hosts, the time-resolved deep sequencing data was used to estimate the growth rate of variants within each host plant and in thrips. The growth rate of each variant can then be used as a proxy for the fitness effect of a variant relative to the reference type. We note, however, that these estimated fitness effects are potentially confounded by variants being linked to other mutations on the same viral genotype/haplotype. Nevertheless, quantifying fitness effects can provide general insights into how selection pressures vary between hosts.

First, the fitness effect of each variant in both *Emilia* and *Datura* was estimated. The joint distribution of fitness effects shows that only a small fraction of variants are estimated to be unconditionally deleterious (Figure 8a). This result is likely due to an ascertainment bias against deleterious variants. Most strongly deleterious mutations were likely excluded since low frequency variants (<3%) were not considered and a variant must persist in the viral population between two or more time points in order to estimate its growth rate (see Methods),. However, there are several mutations that are neutral or beneficial in one host but deleterious in the other, indicative of antagonistic pleiotropy. More surprisingly, there is a strong overall positive correlation between fitness effects across hosts, suggesting that positive rather than antagonistic pleiotropy predominates between plant hosts.

**Figure 8:**
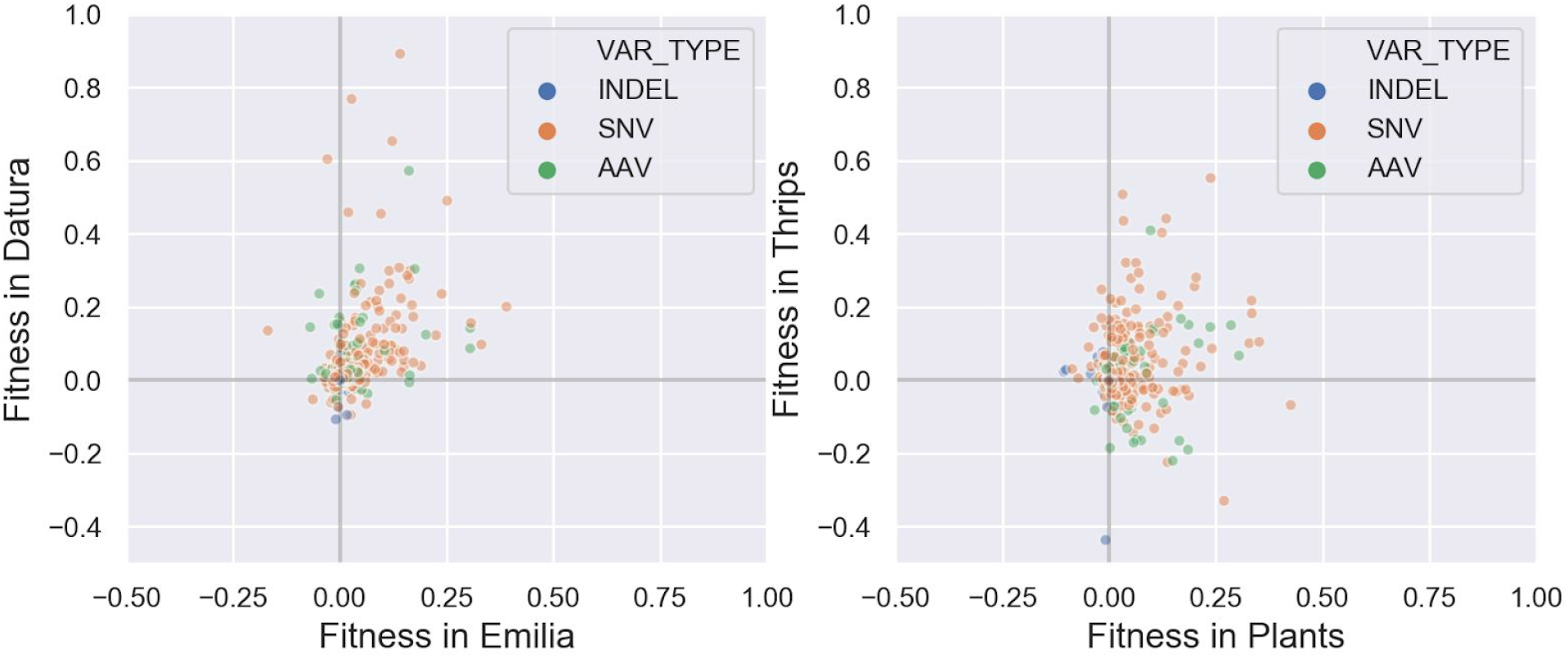
The joint distribution of fitness effects in *Emilia* versus *Datura* **(A)** and in plants versus thrips **(B)**. The fitness effects of individual variants were estimated based on their relative growth rate in each host. Fitness estimates are colored according to the type of mutation: blue = indel; orange = single nucleotide variant; green = amino acid variant.

The joint distribution of fitness effects between plants and thrips is shown in Figure 8b. Fitness effects are positively correlated between plants and thrips, suggesting again that positive pleiotropic effects predominate. However, there are also a rather large number of AAVs that are beneficial in plants but deleterious in thrips, indicating potential fitness conflicts between plants and thrips in certain regions of the genome. To get a better sense of where these fitness conflicts occur, fitness differences between plants and thrips were mapped onto the TSWV genome (Figure 9). Many of the largest fitness differences between plants and thrips are localized on RdRp, and to a lesser extent NSs and G_N_/G_C_. Fitness differences between *Emilia* and *Datura* are distributed over the entire genome, although there are several localized at the N-terminus of NSm.

**Figure 9:**
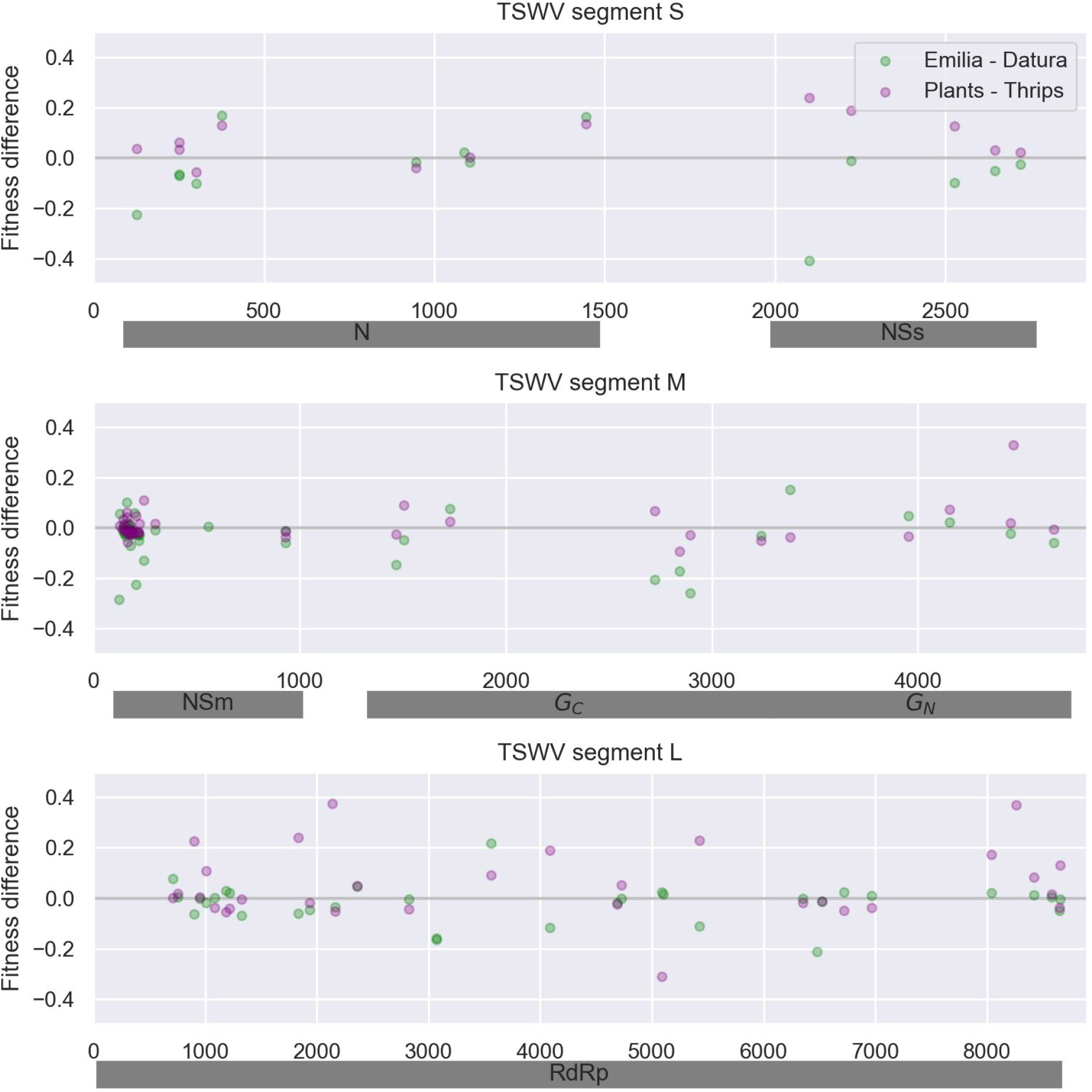
Fitness differences between hosts mapped onto the TSWV genome. Fitness differences between plants and thrips are given as the variant’s fitness in plants minus its fitness in thrips. Positive values therefore indicate greater fitness in plants versus thrips. Likewise, between plant hosts, fitness differences are given as the fitness in *Emilia* minus fitness in *Datura*.

## Discussion

Experimental evolution studies have become a standard approach in virology to investigate how viruses adapt to novel host environments (Turner and Elena, 2000; Duffy *et al.*, 2006; Elena *et al.*, 2008). A large number of these studies have focused on arboviruses or other multi-host pathogens, as virologists have long been interested in how viruses overcome the constraints imposed by alternating between hosts. These experimental evolution studies have often yielded results that challenge long-held assumptions in evolutionary theory. For example, while evolutionary theory largely assumes that performance or fitness tradeoffs will limit simultaneous adaptation to more than one environment, experimental studies have repeatedly demonstrated that viruses can adapt to new hosts with little or no fitness cost in alternate hosts (Weaver *et al.*, 1999; Turner and Elena, 2000; Bedhomme *et al.*, 2012). Likewise, while antagonistic pleiotropy has long been assumed to underlie fitness tradeoffs between environments, recent experimental work has shown that mutations often have positive pleiotropic effects between hosts (Lalic *et al.*, 2011; McGee *et al.*, 2014). In light of this work, we sought to explore how an extreme generalist like TSWV adapts to alternate plant hosts and thrips.

Deep sequencing TSWV populations revealed that much of the genetic diversity present in the initial founder population persisted for multiple passage cycles, with little evidence for bottlenecks in diversity at transmission events. Genetic diversity tended to increase when the virus was passaged through thrips but decrease over the course of a single infection in plants; although this was more evident in *Datura* than *Emilia*. This loss of diversity may be due to the fact that leaves sampled at later time points were more distal from the site of infection. Although TSWV moves systemically through plants, only a subset of the viral population may undergo long-distance transport to new leaves, resulting in distinct founder populations with lower diversity. One might expect to see similar bottlenecks within thrips as the virus must traverse through the midgut to the salivary glands before transmission can occur (Chen *et al.*, 2019). We may have failed to detect bottlenecks in thrips as multiple insects were sampled and then pooled at a single time point to obtain enough RNA for sequencing. However, viral diversity was previously shown to increase in thrips (J. Brown, unpublished), consistent with our results here. Thus, unlike in other plant viruses where vector-borne transmission leads to extreme bottlenecks in viral population sizes and genetic diversity (Gutiérrez *et al.*, 2012), the ability of TSWV to replicate persistently in thrips may largely preserve diversity.

Several amino acid variants rapidly fluctuated in frequency between plant hosts and vectors. The evolutionary dynamics of these variants may provide clues into the selection pressures imposed by different hosts and how the virus adapts to them. In plants, different amino acid variants in the NSm protein were found to be differentially enriched in either *Emilia* and *Datura*. NSm functions as a viral movement protein that is necessary for both short and long distance movement (Lewandowski and Adkins, 2005), and previous reports have implicated NSm in host range determination (Silva *et al.*, 2001). Functional analysis in tobacco plants indicated that amino acid mutations in the N-terminus of NSm abolish tubule formation and cell-to-cell movement, but not long distance movement (Li *et al*, 2009). Interestingly, the first 50 amino acids of the N-terminus are hypervariable at the species level (Silva *et al*, 2001; Li *et al.*, 2009) and hypervariable in our within host populations. Furthermore, we found amino acid variants V17G and N22S are differentially enriched in *Emilia* versus *Datura* (Figure 6), all of which suggests that host-specific changes in NSm may be required for TSWV to move efficiently through different plants.

Several amino acid variants were also found to be enriched in plants versus thrips in a consistent manner across lines. These variants include single amino acid mutations in the silencing suppressor NSs and the glycoprotein G_C_, as well as several in the viral RNA-dependent RNA polymerase (RdRp). As NSs is involved in suppressing RNA silencing in both plants and thrips (Hedil *et al.*, 2015), it is perhaps not surprising that different variants may be favored in plants versus thrips. Interestingly, the amino acid variant E362D in G_C_ appears to be very strongly selected for in thrips but only observed at very low frequencies in plants (Figure 7). Arboviruses have repeatedly been found to adapt to their insect vectors through single amino acid mutations in viral glycoproteins (Brault *et al*, 2004; Tsetsarkin *et al.*, 2007), and in TSWV, G_C_ likely acts as a viral fusion protein that along with G_N_ is essential for transmission in thrips (Sin *et al.*, 2005). But it is less clear why the E362D mutation would be so strongly selected against in plants, since G_N_ and G_C_ are thought to be dispensable in plants (Nagata *et al.*, 2000; Oliver and Whitfield, 2016). Finally, several amino acid variants in RdRp appear to be strongly beneficial in plants but deleterious in thrips. Because the RdRp must replicate and transcribe the viral genome in both plants and thrips, differential selection pressures may arise from the need for RdRp to interact with host-specific cellular factors, which has been shown to be a key determinant of host adaptation in other RNA viruses (Bolvin *et al.*, 2010; Long *et al.*, 2016).

We therefore found some evidence for antagonistic pleiotropy between plants and thrips, and to a lesser extent between *Emilia* and *Datura*, which may place constraints on TSWV’s ability to simultaneously adapt to multiple plant hosts and thrips. Nevertheless, beyond a few sites of apparent conflict in the genome, the fitness effects estimated between hosts largely show that positive rather than antagonistic pleiotropy predominates. It is therefore tempting to speculate that this tendency towards positive pleiotropy endows TSWV with the ability to find beneficial mutations in new hosts without a concomitant loss of fitness in previous hosts, allowing TSWV to rapidly expand its host range. Moreover, even if antagonistic pleiotropy does arise at particular sites in the genome, the ability to maintain extensive genetic diversity between transmission events may allow for variants that are deleterious in the current host to be stored and retrieved in future hosts where that mutation may be beneficial. Thus, the ability to persistently replicate and thereby avoid a narrow transmission bottleneck may allow TSWV to more readily adapt to new hosts than other viruses.

While we were able to estimate the relative fitness effects of variants between hosts, one serious limitation of our study is that the absolute fitness of viral populations between hosts were not directly measured. Furthermore, the present study only considered fitness in alternate plant hosts and not between different thrips species. Like many other plant viruses, TSWV is considered to be a plant host generalist but a vector specialist (Lefeuvre *et al.*, 2019). Indeed, only 9 of more than 7,000 described thrips species are known to be competent vectors of TSWV (Jones *et al.*, 2005; Chen *et al.*, 2019), and particular genetic isolates of TSWV appear to be intimately adapted to local thrips populations (Jacobson and Kennedy, 2013). Future work by our group will therefore look at differences in absolute fitness between hosts and whether it is more difficult for TSWV to adapt to new plant hosts or new vector species.

## Materials & Methods

### Experimental Passaging

A TSWV-infected tomato plant (var. Celebrity) was collected from a field near Apex, North Carolina in August of 2018. The fruit tissue was immediately used for mechanical inoculation onto a 20 day-old *Emilia sonchifolia* plant (referred to as P0 above). Both the fruit and leaf tissue from the same plant were preserved at −80 °C for later RNA extraction and sequencing. The mechanically inoculated *Emilia* plant was used as source material for viral passaging via thrips and maintained under greenhouse conditions within an insect cage.

Western flower thrips (*Frankliniella occidentalis*) were used as the vector species for all passages. Thrips were obtained from a laboratory colony maintained at 27 °C, ca. 55% RH and under continuous light on insecticide-free cabbage (*Brassica oleracea* var*. capitata* L.) foliage in 0.35 L plastic food containers (Fabri-Kal Corp., Kalamazoo, MI) ventilated with thrips-proof screen (81 × 81mesh; Bioquip Products, Inc., Rancho Dominguez, CA). At each transmission cycle, approximately 100 adult females from the colony were confined in a rearing container and allowed to oviposit for 24 hours through a stretched Parafilm^®^ membrane into a 3% sucrose solution contained in a 9 cm Petri dish. Following oviposition, the eggs were collected by filtering the sucrose solution through filter paper and rinsing any eggs attached to the membrane on to the filter paper with distilled water. To obtain viruliferous adults, the filter paper was positioned on top of an excised TSWV-infected leaf from the designated source plant such that the eggs were sandwiched between the filter paper and the abaxial surface of the infected leaf, which was maintained on moistened filter paper in a sealed rearing container at 27 °C. After four days all eggs had hatched and the larvae were shaken onto a non-infected cabbage leaf and reared to adult. Fresh non-infected cabbage tissue was provided as needed.

At each transmission cycle, groups of eight viruliferous adults (3-7 days post-eclosion) were aspirated on to each *Emilia* or *Datura* seedling (three to four-true leaves). Seedlings were grown separately in 296 ml plastic cups (Solo Cup Company, Lake Forest, IL, USA) with a 25 mm diameter fine mesh screen on the bottom. Thrips were contained on the seedlings by inverting a plastic cup with screened bottom over the seedling and sealing it to the cup containing the plant using Parafilm. After approximately 48 hours, each seedling was sprayed with spinetoram (Radiant^®^ SC; Corteva Agriscience, Indianapolis, USA) to kill the thrips. TSWV infected plants were maintained in a growth chamber under a 16-h photoperiod, 27°C and ca. 50% relative humidity for approximately one month after inoculation.

Three separate experimental lines were developed in which the virus was either alternated (Line 1) between plant hosts (*Emilia sonchifolia* and *Datura stramonium*) or maintained on *Emilia* (Line 2) or *Datura* (Line 3). Approximately 21 days after inoculation, tissue was collected from the plant lines and used to initiate the next passaging round by feeding to western flower thrips. At the final passage cycle, the single host lines were passaged back to the alternate plant species.

### Sample Collection

Plant lines were sampled at four time points following virus transmission (at approximately 7, 14, 21, and 28 days post-infection). The time of infection was defined as when the viruliferous thrips were rendered inactive on the host plant. A sterilized 8-mm diameter cork borer was used to collect tissue from the three most recently emerged leaves. Five leaf disks were sampled in total: 2 disks from the two larger leaves and 1 disk from the smallest leaf. Disks from each plant were pooled and immediately frozen at −80 °C for later RNA extraction.

At each transmission cycle, approximately 40 thrips from the cohort of viruliferous adults used to inoculate test plants were collected into a 1.5 ml microcentrifuge tube at the time that transmission was initiated. These thrips were immediately frozen at −80 °C for later RNA extraction.

### Total RNA Extraction

For plant tissue, five 8 mm diameter leaf disks were placed into a 1.5 ml microcentrifuge with three-3mm Pyrex glass beads (Corning). Sample tubes were then placed in liquid nitrogen followed by bead beating on the Silamat S6 (Ivoclar Vivodent) for 20 seconds. Contrastingly, 40 thrips in a 1.5 ml microcentrifuge were placed into liquid nitrogen then ground via motorized pestle.

Following tissue destruction, TRI Reagent (Zymo Research) was immediately added and vortexed on high with 600 ul TRI Reagent added to plant tissue samples, 300 ul to thrips samples. Samples were incubated for 5 minutes in TRI reagent at room temperature before following manufacturer’s protocol for RNA extraction kits. For plant tissue, the Direct-zol RNA MiniPrep Plus kit (Zymo Research) was utilized and RNA resuspended in 60 ul. For thrips, the Direct-zol MicroPrep kit (Zymo Research) with resuspension in 15 ul. RNA quality was assessed via electrophoresis and on a Nanodrop 1000. All RNA samples were stored at −80 °C.

### cDNA synthesis

For synthesis of cDNA from total RNA extracted from plant and thrips tissue samples, approximately 500 ng of total RNA was used for a 10 ul cDNA synthesis with ProtoScript II (NEB). 15 uM of the appropriate strand-specific primer (IDT; Coralville, IA) (Table 1), 10 mM dNTP, total RNA, and sterile water (up to 5ul total volume) were incubated at 65°C for 5 minutes then placed on ice. Next, 2 ul 5X ProtoScript II Buffer, 0.1 M DTT, 4 U RNase Inhibitor, 100 U ProtoScript II RT, and 1.4 ul sterile water were added. The samples were first incubated at 25°C for 5 minutes, 42°C for 1 hour, then 65°C for 20 minutes before storing at −20°C. All passaging samples were duplicated for two independent sequencing replicates beginning at the cDNA step.

**Table 1.**
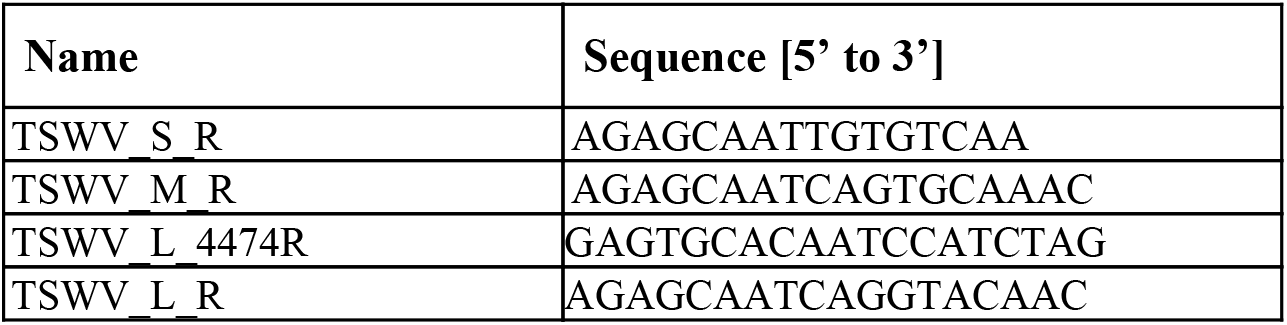
Primers used for segment-specific cDNA synthesis.

### PCR

PCR was used to amplify viral cDNAs to enrich viral representation. 50 ul PCR reactions were set up with approximately 1 ug cDNA and Phusion High-Fidelity DNA Polymerase (NEB) was used. The manufacturer’s protocol was followed and the addition of 1.5 ul DMSO was included. Primers (IDT) utilized were genome segment-specific (Table 2), and the 5’ end included a tail sequence to preferentially bind the Illumina Nextera DNA Flex adapter (to increase terminal end coverage of viral genome segments). PCR reactions were amplified with a BioRad C1000 Thermocycler on the following settings: 98 °C – 30s; 30X: (98 °C – 10s, 52 °C – 30s, 72 °C – 4m); 72 °C – 5m; infinite hold at 12 °C. Expected PCR product sizes were verified via electrophoresis before proceeding to sample purification step.

**Table 2.**
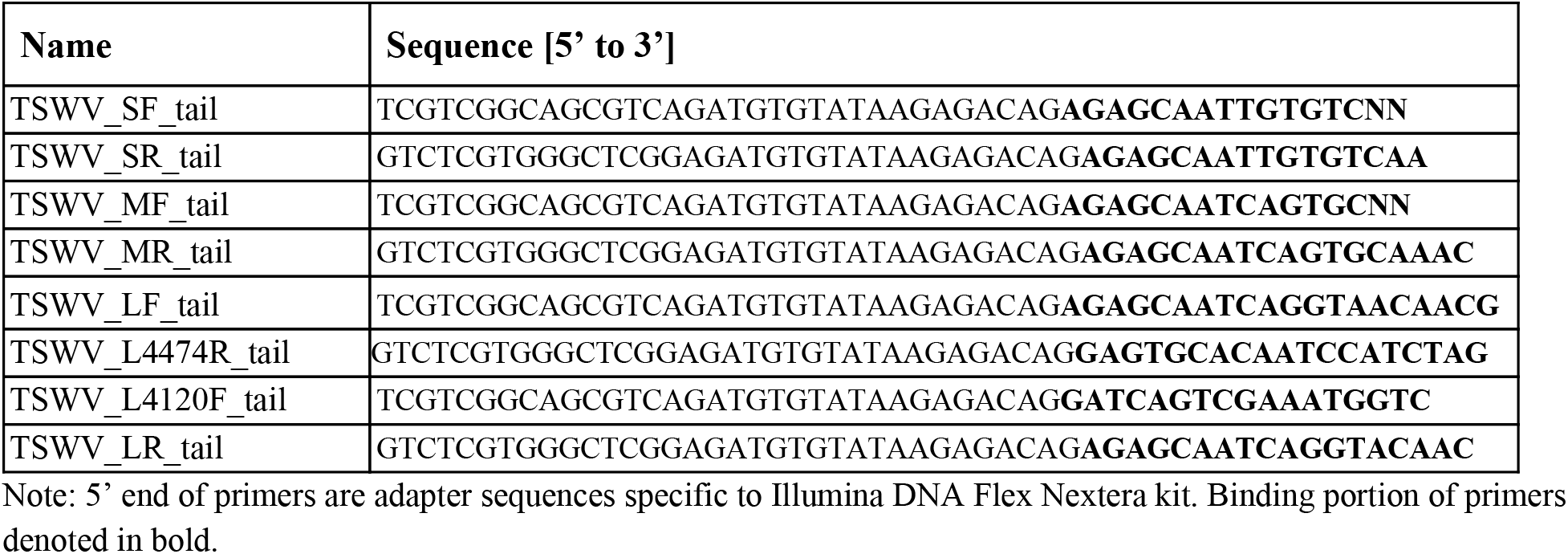
Primers used for segment-specific PCR amplification.

### Sample Purification

Products for the four genome fragments were combined (approximately 200 ul volume when pooled). Combined PCR products were purified with gDNA Clean and Concentrator kit-10 (Zymo Research) via two sequential elutions of 10 ul each (20 ul total volume). Sample concentrations were measured on a Nanodrop 1000.

### Illumina Library Preparation and Sequencing

For deep sequencing, 500 ng of the purified and pooled PCR genome amplicons were prepared for sequencing via Nextera DNA Flex Kit (Illumina, #20018705) with 96 indexes (Illumina, #20018708) according to the manufacturer’s protocol. Library quality was analyzed via Agilent 2200 TapeStation at the NCSU GSL. All sequencing was performed on an Illumina MiSeq instrument at North Carolina State University’s Genomic Science Laboratory to obtain paired end reads of approximately 300 base pairs.

### Sequence analysis

Paired end reads were obtained from two sequencing replicates at each sampling time point from the MiSeq runs. After trimming adapter sequences from the raw reads, sequences were mapped to the TSWV reference genome assembly on GenBank with accession number GCA_000854725.1 (De Haan *et al.*, 1989; 1991) using Bowtie2 (Langmead and Salzberg, 2012). In order to minimize the potential for misalignment due to using a divergent reference genome, we then assembled a new consensus genome sequence from our TF2 field-collected isolate. All paired end reads from our passaging experiments were then aligned against the less divergent TF2 reference genome using the *‘sensitive-local”* preset parameters in Bowtie2. Alignments for each sample were converted into SAM and BAM files for further processing using SAMtools (Li *et al.*, 2009).

To call genetic variants in each viral population, the *mpileup* routine in SAMtools was used to identify single nucleotide variants (SNVs) and indels relative to the TF2 reference in both paired sequencing replicates from each sample. Variants at primer binding sites were first filtered out. The remaining variants were subsequently filtered in ivar using the criteria proposed by the authors (Grubaugh *et al*., 2019). Using their criteria, a variant needed to be present at a frequency of at least 0.03 and obtain an Illumina/Phred quality score of 20 (i.e. a 0.01 sequencing error probability) in both paired sequencing replicates in order to be considered a true variant. Thus, even for sites with a relatively low coverage (<100X), the probability of a variant caused by a sequencing error reaching our threshold frequency of 0.03 is extremely unlikely, with a probability of 10^-6^. Furthermore, while it is possible that an error introduced at the RT or PCR stage could reach a frequency of 0.03, it is extremely unlikely that such an error would occur in both sequencing replicates independently. Our variant calling strategy therefore ensures that all called variants were actually present in the viral population.

From the variants called at each individual sampling time, we created a master list tracking how the frequency of each variant changed over time. We also categorized SNVs as either amino acid variants (AAVs) based on whether their translated sequence was predicted to cause a nonsynonymous substitution in the reference sequence.

### Global TSWV diversity

Within-host genetic diversity was compared to species-level diversity among a global collection of TSWV isolates sampled from different hosts. For this analysis, the same set of sequences as Lian *et al*. (2013) was used which included 53 S, 57 M and 17 L full-length segment sequences. To this collection, we added 23 L segment sequences that have been deposited in GenBank since 2013. GenBank accession numbers for all sequences are provided in Supp. Table 1.

### Estimating fitness effects

The fitness effect of each variant was estimated based on changes in variant frequencies over time within hosts. Following the strategy of Illingworth *et al*. (2014), it is assumed that each variant’s frequency changes over time according to a model of deterministic exponential growth:

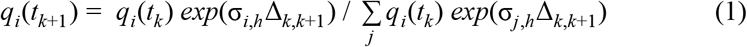

Here, *q_i_*(*t*_*k*+1_) is the predicted frequency of variant *i* at time *t*_*k+1*_ given it’s observed frequency *q*(*t_k_*) at time *t_k_*. The term Δ_*k,k*+1_ is the time elapsed between a pair of sequential samples taken at times *k* and *t*_*k*+1_. The host-specific growth rate of variant *i* in host *h* is given by σ_i,h_. Note that the growth rate of each variant is estimated relative to the reference type since absolute growth rates cannot be estimated because only variant frequencies are observed through time.

The growth rate σ_i,h_ therefore reflects variant *i*’s relative fitness in a particular host, and we seek to estimate these values from observed frequency changes over time. Let *n*_*k+1*_ be a vector holding the number of observed sequence reads representing each variant at time *t_k+1_*. Given the expected variant frequencies *q*(*t*_*k*+1_) predicted under the exponential growth model, we compute the likelihood of observing *n_+1_* assuming a multinomial sampling process:

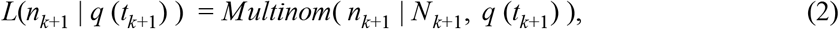

where 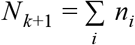, the total depth of coverage at the site of variant *i*.

To obtain a maximum likelihood estimate for the fitness effects, we can then find the value of σ_i,h_ that maximizes the product of the individual multinomial likelihood terms (eq. 2) across all pairs of time points *k* and *k+1* for which we have observed variant frequencies, using eq. 1 above to compute *q_i_*(*t*_*k*+1_) whenever we need to evaluate the likelihood function. We exclude all pairs of time points where the initial frequency of the variant or reference allele was zero at time *t_k_* because in this case eq. 2 is not defined. We also exclude all pairs of time points where *N_k+1_* <100 to minimize variability in our estimates due to a low total depth of coverage at a given sampled time point. Maximum likelihood estimates were obtained by numerically optimizing the likelihood with SciPy’s *minimize* function using Sequential Least Squares Programming.

## Acknowledgments

The authors would like to thank Dr. Tim Sit at NC State for generously providing lab space.

## Data and code availability

Python scripts automating the alignment, variant calling and fitness estimation procedures are available at https://github.com/davidrasm/DeepTSWVSeq. A master list providing the frequency of each variant through time in all three lines is also available in this repository. Raw sequence reads have been submitted to NCBI’s Sequence Read Archive as a single BioProject.

---

**Supp. Figure 1:**
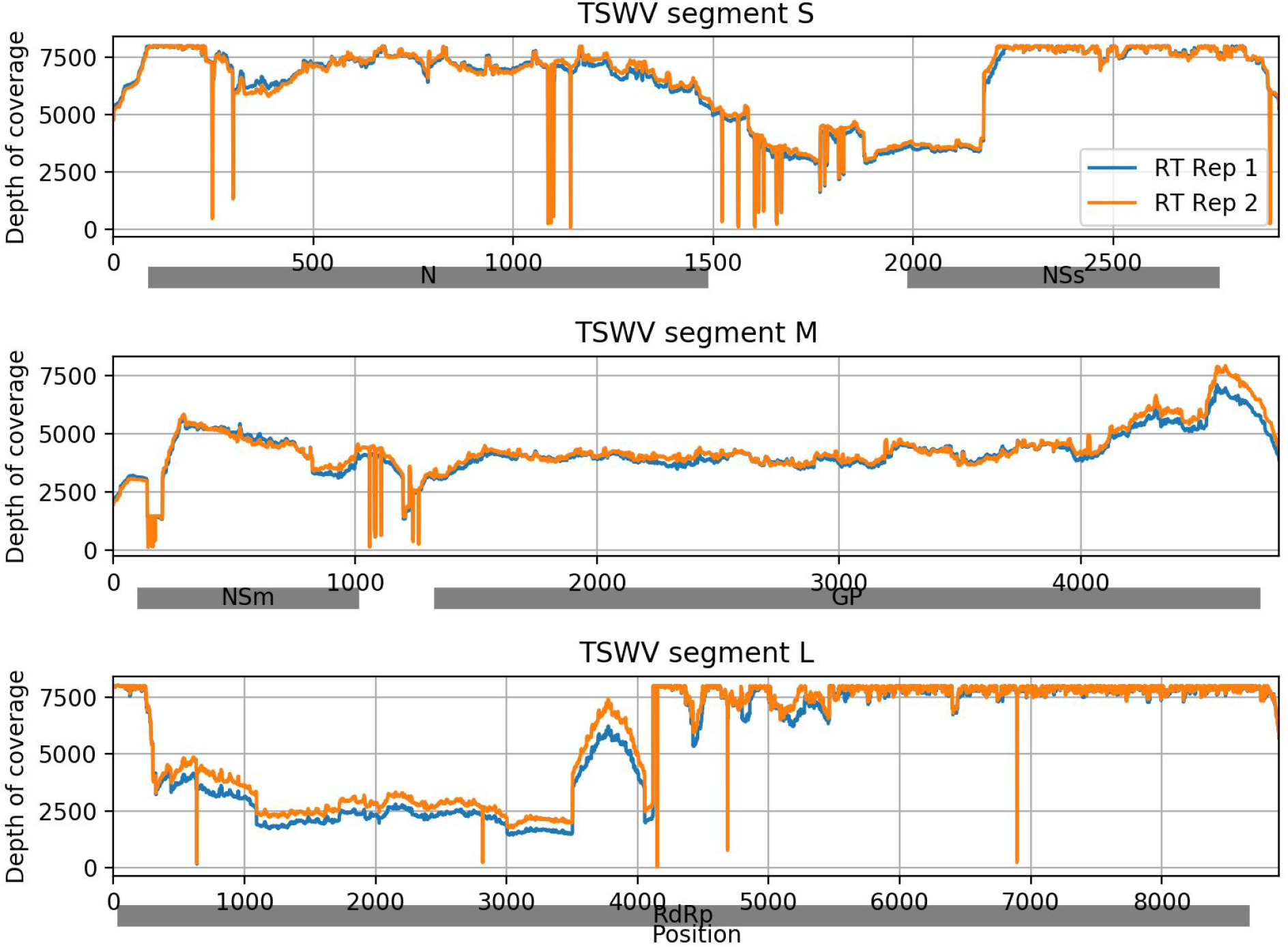
Sequencing coverage across the TSWV genome for the viral population at passage P0. Blue and orange lines show coverage obtained in paired sequence replicates obtained from independent reverse transcription reactions.

**Supp. Figure 2:**
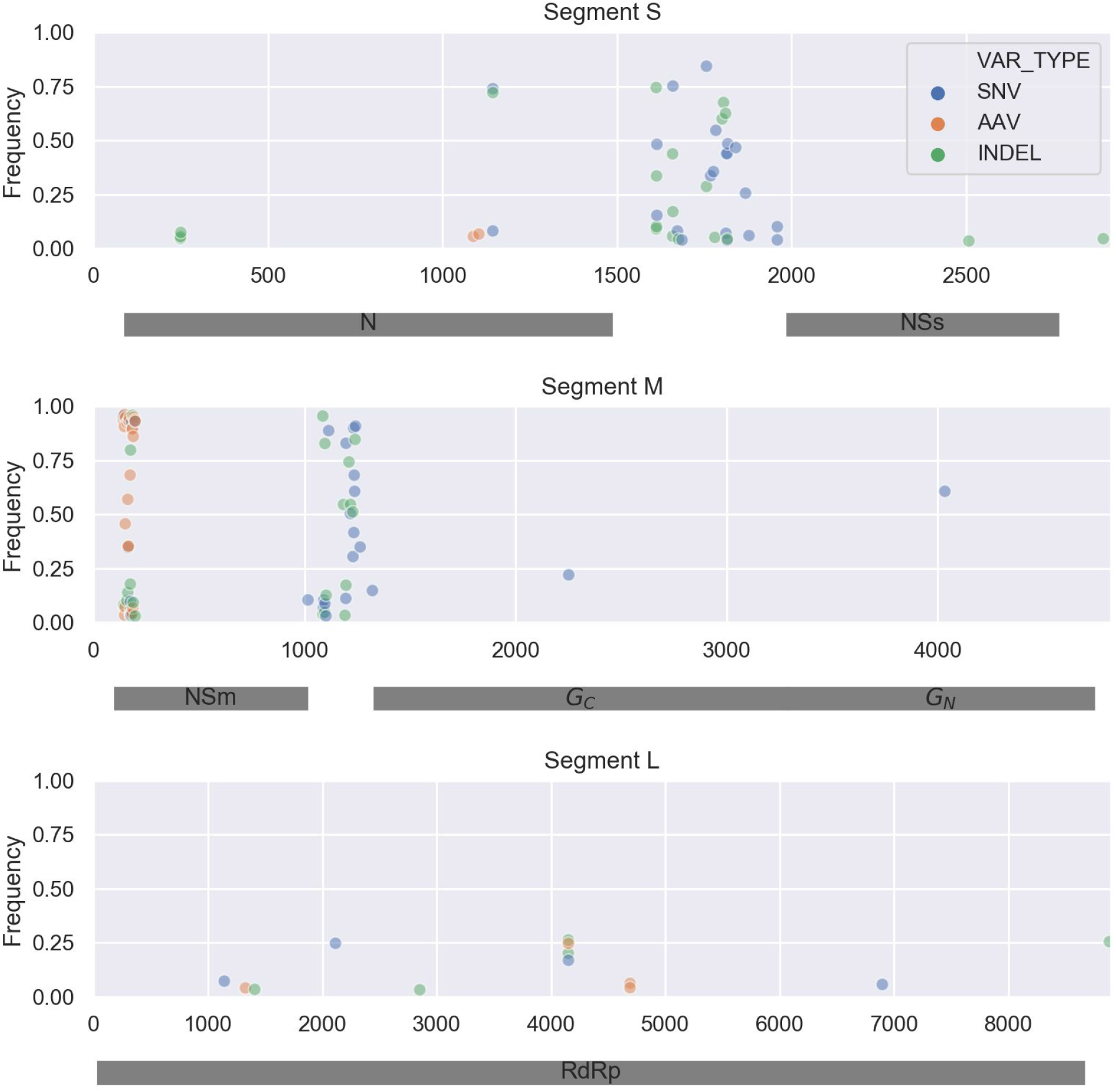
Viral genetic diversity in a naturally infected tomato leaf (TL2) collected in the field from the same plant as TF2. The frequency of each single nucleotide variant (SNV), amino acid variant (AAV) and indel is plotted at its respective position along the three segments of TSWV’s genome. Shaded grey rectangles represent protein coding regions. Note: this is TL2_allVarFreqs_noLabels.png

**Supp. Figure 3:**
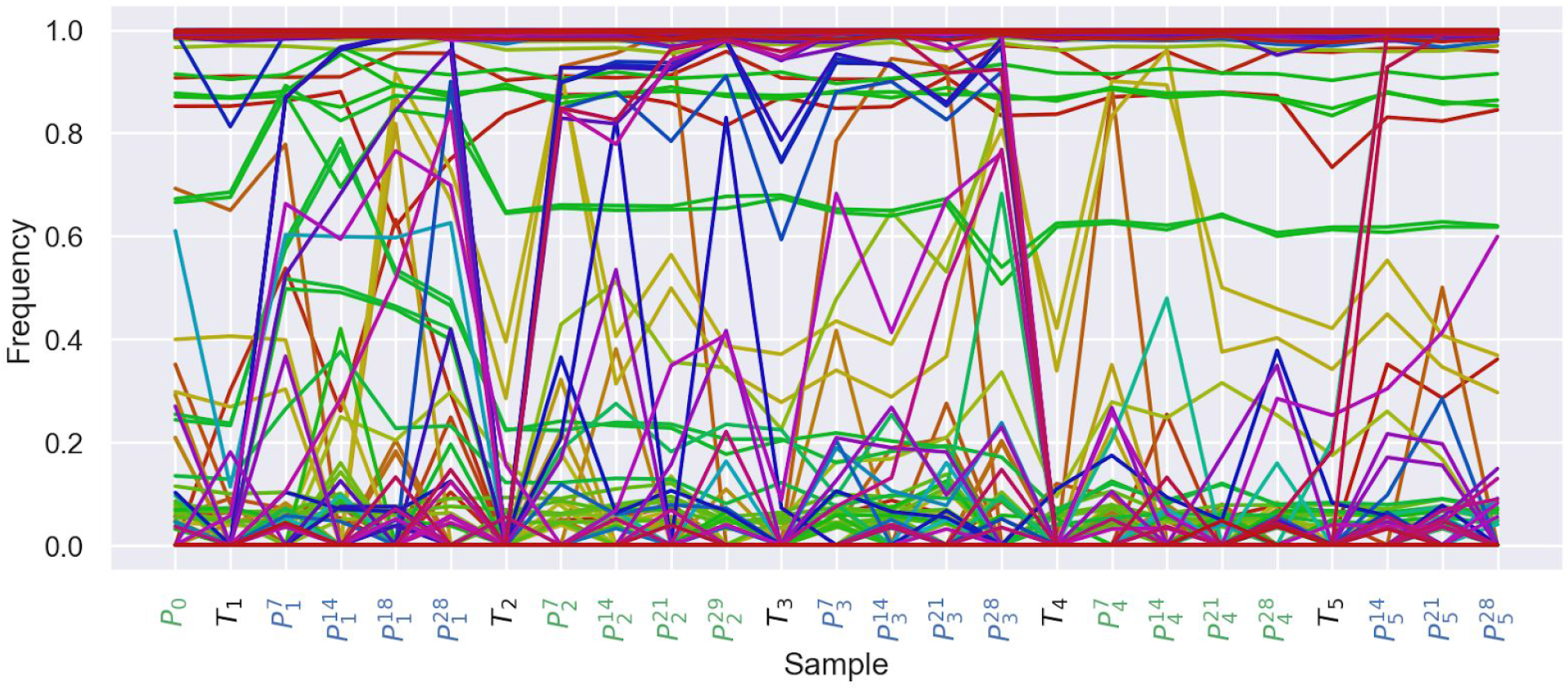
Time series showing the evolutionary dynamics of all single nucleotide variants through time in the *Alternating* Line. Sampling time points are colored by host; green = *Emilia*, black = thrips and blue = Datura.

